# Individual differences in motor noise and adaptation rate are optimally related

**DOI:** 10.1101/238865

**Authors:** Rick van der Vliet, Maarten A. Frens, Linda de Vreede, Zeb D. Jonker, Gerard M. Ribbers, Ruud W. Selles, Jos N. van der Geest, Opher Donchin

**Affiliations:** Departments of Neuroscience; Departments of Rehabilitation Medicine, Erasmus MC, 3015 CN, Rotterdam, The Netherlands; Departments of Erasmus University College, 3011 HP, Rotterdam, The Netherlands; Departments of Rijndam Rehabilitation Centre, 3015 LJ, Rotterdam, The Netherlands; Department of Plastic and Reconstructive Surgery, Erasmus MC, 3015 CN, Rotterdam, The Netherlands; Department of Biomedical Engineering and Zlotowski Center for Neuroscience, Ben Gurion University of the Negev, 8499000, Be’er Sheva, Israel

## Abstract

Individual variations in motor adaptation rate were recently shown to correlate with movement variability or “motor noise” in a forcefield adaptation task. However, this finding could not be replicated in a meta-analysis of visuomotor adaptation experiments. Possibly, this inconsistency stems from noise being composed of distinct components which relate to adaptation rate in different ways. Indeed, previous modeling and electrophysiological studies have suggested that motor noise can be factored into planning noise, originating from the brain, and execution noise, stemming from the periphery. Were the motor system optimally tuned to these noise sources, planning noise would correlate positively with adaptation rate and execution noise would correlate negatively with adaptation rate, a phenomenon familiar in Kalman filters. To test this prediction, we performed a visuomotor adaptation experiment in 69 subjects. Using a novel Bayesian fitting procedure, we succeeded in applying the well-established state-space model of adaptation to individual data. We found that adaptation rate correlates positively with planning noise (r=0.27; 95%HDI=[0.05 0.50]) and negatively with execution noise (r=−0.41; 95%HDI=[−0.63 −0.16]). In addition, the steady-state Kalman gain calculated from state and execution noise correlated positively with adaptation rate (r = 0.31; 95%HDI = [0.09 0.54]). These results suggest that motor adaptation is tuned to approximate optimal learning, consistent with the “optimal control” framework that has been used to explain motor control. Since motor adaptation is thought to be a largely cerebellar process, the results further suggest the sensitivity of the cerebellum to both planning noise and execution noise.

**SIGNIFICANCE STATEMENT:** Our study shows that the adaptation rate is optimally tuned to planning noise and execution noise across individuals. This suggests that motor adaptation is tuned to approximate optimal learning, consistent with “optimal control” approaches to understanding the motor system. In addition, our results imply sensitivity of the cerebellum to both planning noise and execution noise, an idea not previously considered. Finally, our Bayesian statistical approach represents a powerful, novel method for fitting the well-established state-space models that could have an influence on the methodology of the field.

## INTRODUCTION

As children we all learned: some of us move with effortless grace and others are frankly clumsy. Underlying these differences are natural variations in acquiring, calibrating and executing motor skill, which have been related to genetic (Frank et al., 2009; Fritsch et al., 2010; McHughen et al., 2010) and structural factors (Tomassini et al., 2011). Recently, it has been suggested that differences between individuals in the rate of motor adaptation (i.e. the component of motor learning responsible for calibrating acquired motor skills to changes in the body or environment (Shadmehr et al., 2010)), correlate with movement variability, or motor noise (Wu et al., 2014). However, this finding was not supported by a recent meta-analysis of adaptation experiments (He et al., 2016). This inconsistency may arise because motor noise has multiple components with differing relation to adaptation rate. Our study characterizes the relationship between adaptation rate and motor noise and suggests that adaptation rate varies optimally between individuals in the face of multiple sources of motor variability.

Motor noise has many physiological sources such as motor preparation noise in (pre)motor networks, motor execution noise, and afferent sensory noise (Faisal et al., 2008). Modeling (Cheng and Sabes, 2006, 2007; van Beers, 2009) and physiological studies (Churchland et al., 2006; Chaisanguanthum et al., 2014) have divided the multiple sources of motor noise into planning noise and execution noise (see Figure 1A). Planning noise is believed to arise from variability in the neuronal processing of sensory information, as well as computations underlying adaptation and maintenance of the states in time (Cheng and Sabes, 2007). Indeed, electrophysiological studies in macaques show that activity in (pre)motor areas of the brain is correlated with behavioral movement variability (Churchland et al., 2006; Chaisanguanthum et al., 2014). Similar results have also been seen in humans using fMRI (Haar et al., 2017). In contrast, execution noise apparently originates in the sensorimotor pathway. In the motor pathway, noise stems from the recruitment of motor units (Harris and Wolpert, 1998; Jones et al., 2002; van Beers et al., 2004). Motor noise is believed to dominate complex reaching movements with reliable visual information (van Beers et al., 2004). In addition, sensory noise stems from the physical limits of the sensory organs and has been proposed to dictate comparably simpler smooth pursuit eye movements (Bialek, 1987; Osborne et al., 2005). Planning and execution noise might affect motor adaptation rate in different ways.

**Figure 1.**
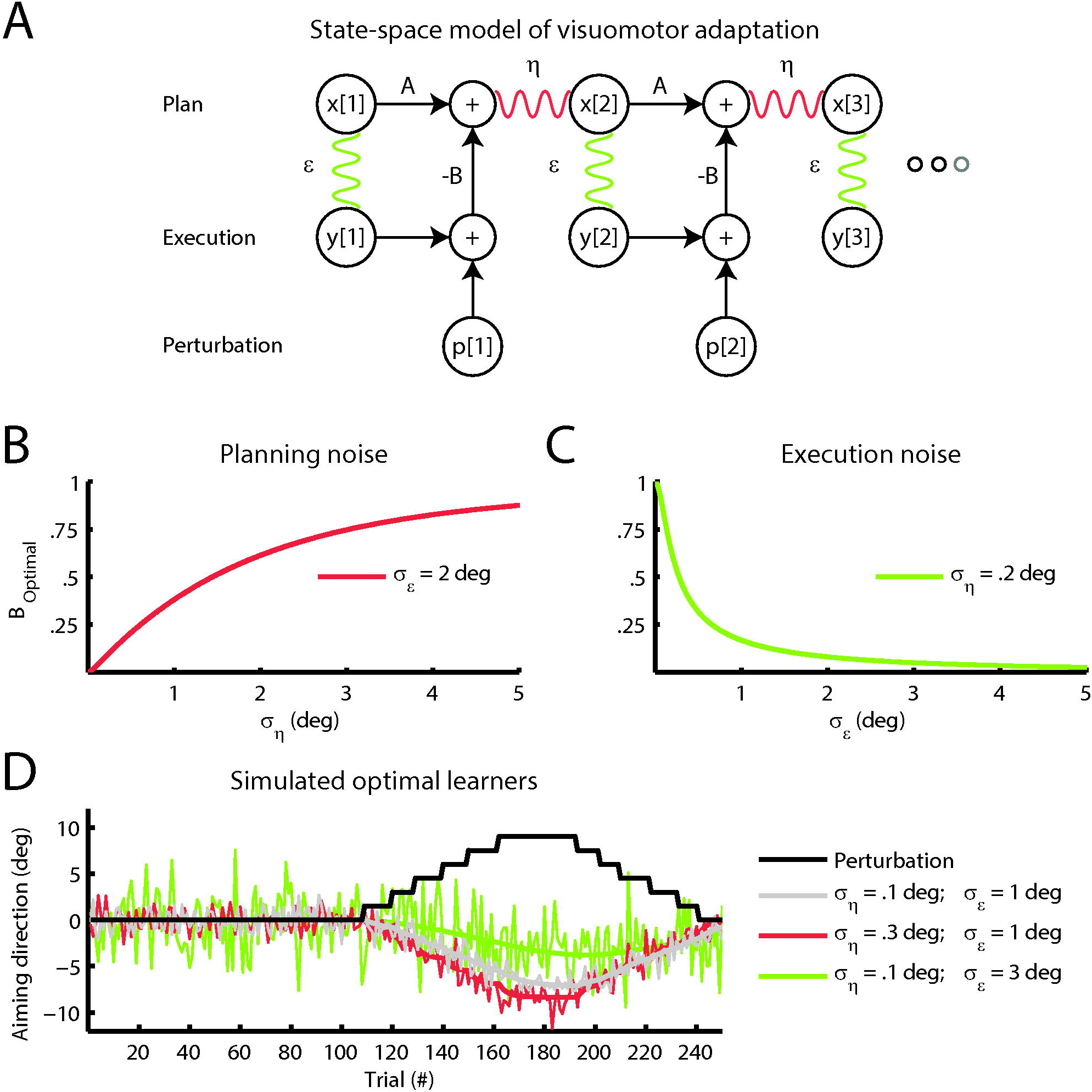
State and execution noise have opposing effects on visuomotor adaptation. **A.** State-space model of visuomotor adaptation. Aiming directions are planned on trial *x*[2] as a linear combination of the state on the previous trial *x*[1] multiplied by a retentive factor *A* minus the error *e*[1] on the previous trial multiplied with learning factor *B*. In addition, the movement plan is distorted by the random process *η*. The actual aiming direction *y*[2]is the planned movement distorted by the random process *ϵ*. The error *e*[1] is the sum of the aiming direction relative to the target *y*[1] and external perturbation *p*[1]. **B.** Planning noise and optimal adaptation rate *B_Optimal_* (defined as the Kalman gain). The optimal adaptation rate increases with planning noise *σ_η_*. In this figure, *σ_ϵ_* was kept constant at 2°. **C.** Execution noise and optimal adaptation rate *B_Optimal_* (defined as the Kalman gain). The optimal adaptation rate decreases with execution noise *σ_ϵ_*. In this figure, *σ_η_* was kept constant at 0.2°. **D.** Simulated optimal learners. At trial 110, a perturbation (black line) is introduced that requires the optimal learners to adapt their movement. The gray learner has low planning noise *σ_η_* = 0.1° and execution noise *σ_ϵ_* = 1°. The red learner has a higher planning noise *σ_η_* = 0.3° than the gray learner *σ_η_* = 0.1°. This causes the red learner to adapt faster. The green learner has a higher execution noise than the gray learner *σ_ϵ_* = 3°. This causes the green learner to adapt more slowly. For all learners, the thick line shows the average, thin line a single noisy realization.

Motor adaptation has long been suspected to be sensitive to planning noise and execution noise. Models of adaptation incorporating both planning and execution noise have been shown to provide a better account of learning than single noise models (Cheng and Sabes, 2006, 2007; van Beers, 2009). In addition, manipulating the sensory reliability by blurring the error feedback, effectively increasing the execution noise, can lower the adaptation rate (Baddeley et al., 2003; Burge et al., 2008; Wei and Körding, 2010; van Beers, 2012) whereas manipulating state estimation uncertainty by temporarily withholding error feedback, effectively increasing the planning noise, can elevate the adaptation rate (Wei and Körding, 2010). These studies not only suggest that adaptation rate is tuned to multiple sources of noise, but also indicate that this tuning process is optimal and can therefore be likened to a Kalman filter (Kalman, 1960). Possibly, differences in adaptation rate between individuals correlate with planning noise and execution noise according to the same principle, predicting faster adaptation for people with more planning noise and slower adaptation for people with more execution noise (He et al., 2016) (Figure 1C and Figure 1D).

To test the relation between adaptation rate and planning noise and execution noise across individuals, we performed a visuomotor adaptation experiment in 69 healthy subjects. We fitted a state-space model of trial-to-trial behavior (Cheng and Sabes, 2006, 2007) using Bayesian statistics to extract planning noise, execution noise and adaptation rate for each subject. We show that the adaptation rate is sensitive to both types of noise and that this sensitivity matches predictions based on Kalman filter theory.

## METHODS

### Subjects

We included 69 right-handed subjects between October 2016 and December 2016, without any medical conditions that might interfere with motor performance (14 men and 55 women; age M=21 years, range 18 - 35 years; handedness score M=79; range 45 – 100). Subjects were recruited from the Erasmus MC University Medical Centre and received a small financial compensation. The study was performed in accordance with the Declaration of Helsinki and approved by the medical ethics committee of the Erasmus MC University Medical Centre.

### Experimental procedure

#### Visuomotor adaptation

Subjects were seated in front of a horizontal projection screen while holding a robotic handle in their dominant right hand (previously described in (Donchin et al., 2012)). The projection screen displayed the location of the robotic handle (“the cursor”; yellow circle 5 mm radius), start location of the movement (“the origin”, white circle 5 mm radius), and target location of the movement (“the target”, white circle 5 mm radius) on a black background (see Figure 2A). Position of the origin on the screen was fixed throughout the experiment, approximately 40 cm in front of the subject at elbow height, while the target was placed 10 cm from the origin at an angle of −45°, 0°or 45°. To remove direct visual feedback of hand position, subjects wore an apron that was attached to the projection screen around their neck.

**Figure 2.**
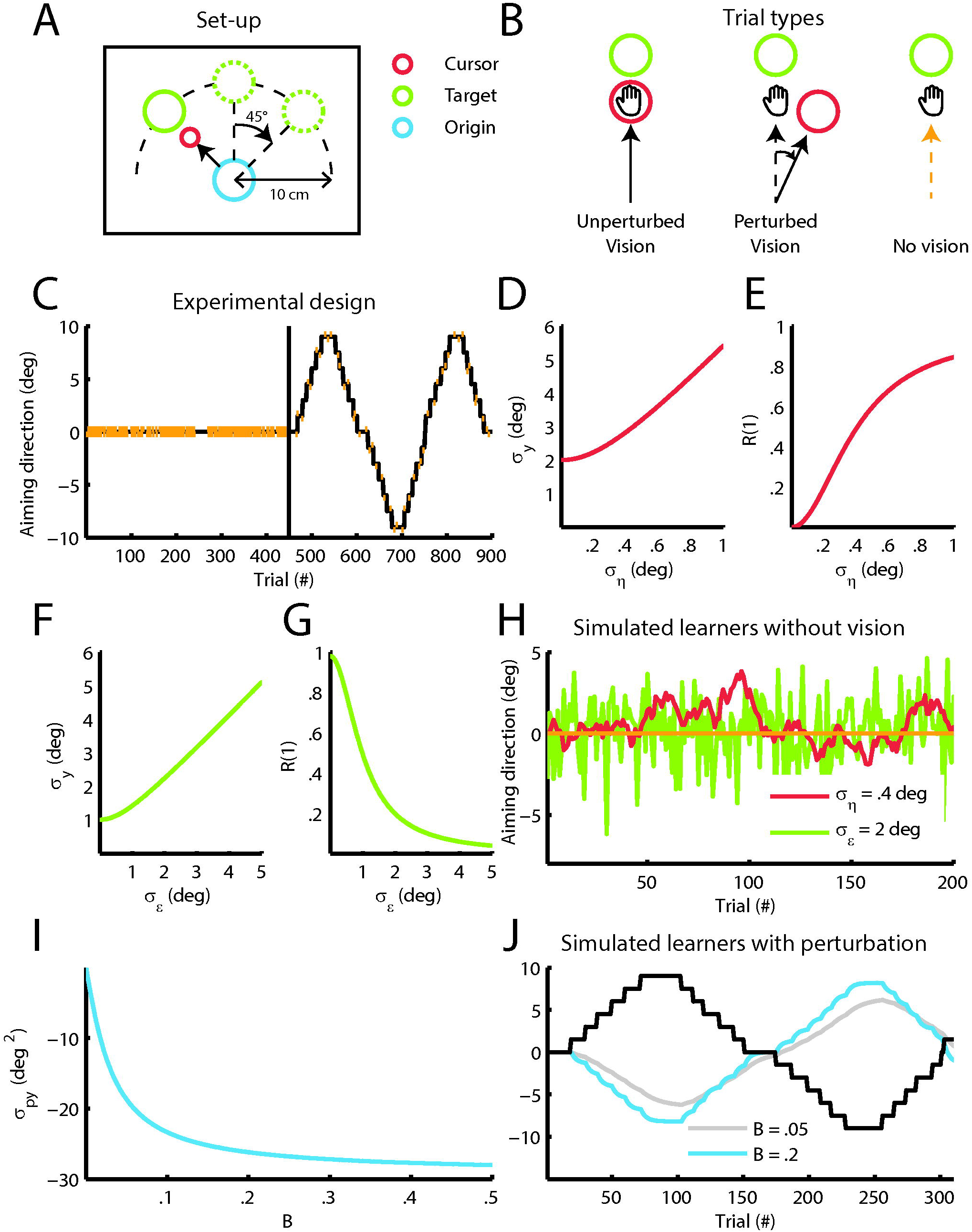
Measurements of state and execution noise and adaptation rate in a visuomotor adaptation experiment. **A.** Set-up. The projection screen displayed the location of the robotic handle (“the cursor”), start location of the movement (“the origin”), and target of the movement (“the target”) on a black background. The position of the origin on the screen was fixed throughout the experiment, while the target was placed 10 cm from the origin at an angle of −45°, 0° or 45°. **B.** Trial types. The experiment included vision unperturbed and perturbed trials and no vision trials. In vision unperturbed trials, the cursor was shown at the position of the handle during the movement. The cursor was also visible in vision perturbed trials but at a predefined angle from the vector connecting the origin and the handle. In no vision trials, the cursor was turned off when movement onset was detected and therefore only visible at the start of movement to help subjects keep the cursor at the origin. **C.** Experimental design. The baseline block consisted of 225 vision unperturbed trials and 225 no vision trials (indicated by vertical red lines). The perturbation block had 50 no vision trials and 400 vision trials, with every block of nine trials containing one no vision trial. Most vision trials were perturbed vision trials whose perturbation magnitudes formed a staircase running from -9 to 9°. **D.** Simulation of planning noise *σ_η_* and aiming direction standard deviation *σ_y_*. *σ_y_* increases with *σ_η_* (calculated for *A* = 0.98, *B* = 0, *σ_ϵ_* = 2°). **E.** Simulation of planning noise *σ_η_* and aiming direction lag-1 autocorrelation *R*(1). *R*(1) increases with *σ_η_* (calculated for *A* = 0.98, *B* = 0, *σ_ϵ_* = 2°). **F.** Simulation of execution noise *σ_ϵ_* and aiming direction standard deviation *σ_y_. σ_y_* increases with *σ_ϵ_* (calculated for *A* = 0.98, *B* = 0, *σ_η_* = 0.2°). **G.** Simulation of execution noise *σ_ϵ_* and aiming direction lag-1 autocorrelation *R*(1). *R*(1) decreases with *σ_ϵ_* (calculated for *A* = 0.98, *B* = 0, *σ_η_* = 0.2°). **H.** Simulated learners without vision. The green and red traces show a single realization of two learners with either high planning noise (red learner *σ_η_* = 0.4° and *σ_ϵ_* = 0°) or high execution noise (green learner *σ_η_* = 0° and *σ_ϵ_* = 2°). Both sources increase the aiming noise, but planning noise leads to correlated noise whereas execution noise leads to uncorrelated noise. This property can be seen from the relation between sequential trials. For the red learner sequential trials are often in the same (positive or negative) direction. For the green learner sequential trials are in random directions. This is captured by the lag-1 autocorrelation. **I.** Simulation of *σ_py_* between the perturbation *p* and aiming direction *y*, and adaptation rate *B. σ_py_* gets more negative for increasing *B* (simulated with *A* = 0.98). **J.** Simulated learners with perturbation. The gray and blue lines show a simulated slow (*A* = 0.98, *B* = 0.05) and fast learner (*A* = 0.98, *B* = 0.2). The fast learner tracks the perturbation signal more closely than the slow learner. This property is captured by the covariance between the perturbation and the aiming direction.

Subjects were instructed to make straight shooting movements from the origin towards the target and to decelerate only when they passed the target. A trial ended when the distance between the origin and cursor was at least 10 cm or when trial duration exceeded 2 seconds. At this point, movements were damped with a force cushion (damper constant 3.5 Ns/m, spring constant 35 N/m) and the cursor was displayed at its last position until the start of the next trial to provide position error feedback. Furthermore, velocity feedback was given to keep movement velocity in a tight range. The target dot turned blue if movement time on a particular trial was too long (>600 ms), red if movement time was too short (<200 ms) and remained white if movement time was in the correct time range (200-600 ms). During presentation of position and velocity feedback, the robot pushed the handle back to the starting position. Forces were turned off when the handle was within 0.5 cm from the origin. Concurrently, the cursor was projected at the position of the handle again and subjects had to keep the cursor within 0.5 cm from the origin for 1 second to start the next trial.

The experiment included vision unperturbed, vision perturbed and no vision trials (see Figure 2B). In vision unperturbed trials, the cursor was shown at the position of the handle during the movement. The cursor was also visible in vision perturbed trials but at a predefined angle from the vector connecting the origin and the handle. In no vision trials, the cursor was turned off when movement onset was detected (see below) and was visible only at the start of the trial to help subjects keep the cursor at the origin.

The entire experiment lasted 900 trials with all three target directions (angle of −45°, 0° or 45°) occurring 300 times in random order. The three different trial types were used to build a baseline and a perturbation block (see Figure 2C). We designed the baseline block to estimate planning noise and execution noise, and obtain variance statistics (standard deviation and lag-1 autocorrelation) related to these noise parameters. Therefore, we limited trial-to-trial adaptation by including a large number of no vision trials (225 no vision trials) as well as vision unperturbed trials (225 vision unperturbed trials). The order of the vision unperturbed trials and no vision trials was randomized except for trials 181-210 (no vision trials) and trials 241-270 (vision unperturbed trials). We designed the perturbation block to estimate trial-to-trial adaptation, and obtain variance statistics related to trial-to-trial adaptation (covariance between perturbation and aiming direction). Therefore, the perturbation block consisted of a large number of vision trials (400 vision trials) and a small number of no vision trials (50 no vision trials), with every block of nine trials containing one no vision trial. Every eight to twelve trials, the cursor perturbation size changed with an incremental 1.5° step. These steps started in the positive direction until reaching 9° and then switched sign to continue in the opposite direction until reaching −9°. This way, a perturbation signal was constructed with three “staircases” lasting 150 trials each (see Figure 2C). The experiment was briefly paused every 150 trials.

#### Data Collection

The experiment was controlled by a C++ program developed in-house. Position and velocity of the robot handle were recorded continuously at a rate of 500 Hz. Velocity data was smoothed with an exponential moving average filter (smoothing factor=0.18s). Trials were analyzed from movement start (defined as the time point when movement velocity exceeds 0.03 m/s) to movement end (defined as the time point when the distance from the origin is equal to or larger than 9.5 cm). Aiming direction was defined as the signed (+ or −) angle in degrees between the vector connecting origin and target and the vector connecting movement start and movement end including the visual perturbation. The clockwise direction was defined positive. Peak velocity was found by taking the maximum velocity in the trial interval. We calculated peak velocity to investigate its relationship with planning noise and execution noise. Trials with (1) a maximal displacement below 9.5 cm, (2) an aiming direction larger than 30° or (3) a duration longer than 2 seconds were removed from further analysis (2% of data).

#### Visuomotor adaptation model

The aiming direction was modeled with the following state-space equation (see Figure 1A) (Cheng and Sabes, 2006, 2007):

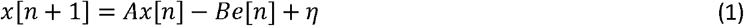

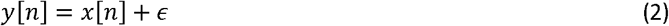

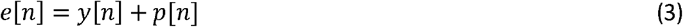

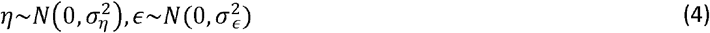

In this model, *x*[*n*] is the aiming plan and *y*[*n*] the executed aiming movement. Error e[*n*] on a particular trial is the sum of *y*[*n*] and the perturbation *p*[*n*]. The learning terms are *A*, which represents retention of the aiming plan over trials, and *B*, the fractional change from error *e*[*n*], The aiming plan is affected by planning noise process *η*, modeled as a zero-mean Gaussian with standard deviation *σ_η_*, and execution noise process e, modeled as a zero-mean Gaussian with standard deviation *σ_ϵ_*.

### Statistics

We designed a statistical procedure to fit the state-space model described in equations (1)–(4) to the data of individual subjects using Markov-chain Monte-Carlo sampling (Kruschke, 2010) implemented in OpenBugs (ver 3.2.3, OpenBugs Foundation available from: http://www.openbugs.net/w/Downloads) with three 50,000 samples chains and 20,000 burn-in samples. A single estimate per subject *s* was made for *A*[*s*] and *B*[*s*] using all trials. Separate estimates were made per subject in the baseline and perturbation block for *B_Block_*[*s*], 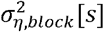 and 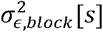. Our main analysis focused on the relation between *B_Perturbation_*[*s*], *σ_η,baseline_*[*s*] and *σ_ϵ,baseline_*[*s*] (see below), similar to (Wu et al., 2014).

We defined a logistic normal distribution as a prior for *A*[*s*] and *B_Block_*[*s*], a normal distribution as a prior for *B*[*s*] and an inverse gamma distribution as a prior for 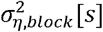 and 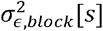:

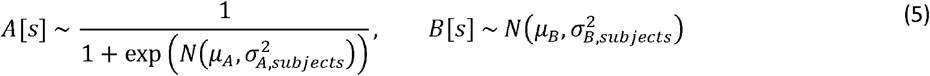

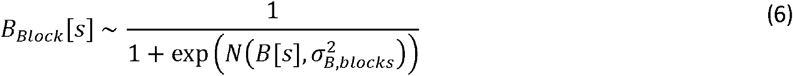

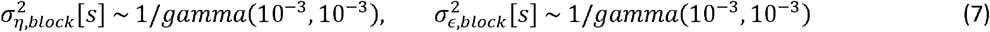

Priors for *μ_A_* and *μ_B_* were selected from a normal distribution and priors for 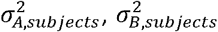 and 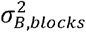 from a gamma distribution:

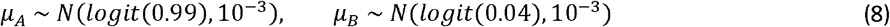

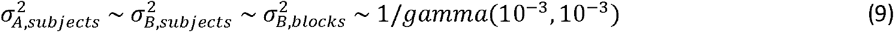

The mode of the samples per parameter and subject was used for further calculations.

We calculated normalized Bayesian linear regression coefficients to investigate the relation between adaptation rate *B_Perturbation_* and noise terms *σ_η,baseline_* and *σ_ϵ,baseline_* (Openbugs, three 50,000 samples chains and 20,000 burn-in samples). The dependent variable was modeled as a t-distribution with the regression model as the mean. As priors, we used uniform distributions (range −1 to +1) for the coefficients, normal distributions for the intercepts (zero mean, precision 10^−6^), gamma distributions for the model error (shape and rate parameter 10^−3^) and a shifted exponential prior (rate parameter 1/29) on the degrees of freedom (Kruschke, 2010). This way, we evaluated (1) an intercept model, and (2) an intercept with planning noise *σ_η,block_* and execution noise *σ_ϵ,block_* model. Model quality was determined by calculating the difference in the deviance information criterion (DIC) between that model and the intercept model (Δ*DIC* = *DlC_Model_* − *DlC_Intercept Model_*). The DIC assigns a score to a model by penalizing the complexity and rewarding the fit. Better models have lower DICs and better models therefore have a negative Δ*DIC*. In addition, we tested correlations between parameters with Bayesian Pearson correlation coefficients, using similar priors as for the linear regression.

Statistical results are reported as the mode of the effect size with 95% highest density intervals (HDIs). Model estimates are plotted as the mode with 68% HDIs, similar to the standard deviation interval.

## RESULTS

### Modelling learning and noise in visuomotor adaptation

We designed a visuomotor adaptation task to capture baseline variability in a baseline block and adaptation to a perturbation in a perturbation block (Tseng et al., 2007) (see Figure 2A-C). We fitted the state-space model described in equations (1)–(4) to the data of individual subjects using Markov-chain Monte-Carlo sampling. A single estimate per subject *s* was made for *A*[*s*] and *B*[*s*] using all trials. Separate estimates were made per subject in the baseline and perturbation block for *B_Block_*[*s*], *σ_η,block_*[*s*] and *σ_ϵ,block_*[*s*] (similar to (Wu et al., 2014)). Our main analysis was the regression of *B_Perturbation_*[*s*] onto *σ_η,baseline_*[*s*] and *σ_ϵ,baseline_*[*s*].

Standard deviation of aiming direction calculated across the 69 subjects illustrates the differences in movement behavior between people (Figure 3A). The state-space model which captures retention *A*, learning from error *B_Block_*, planning noise *σ_η,block_* and execution noise *σ_ϵ,block_* shows good agreement with the average aiming direction.

**Figure 3.**
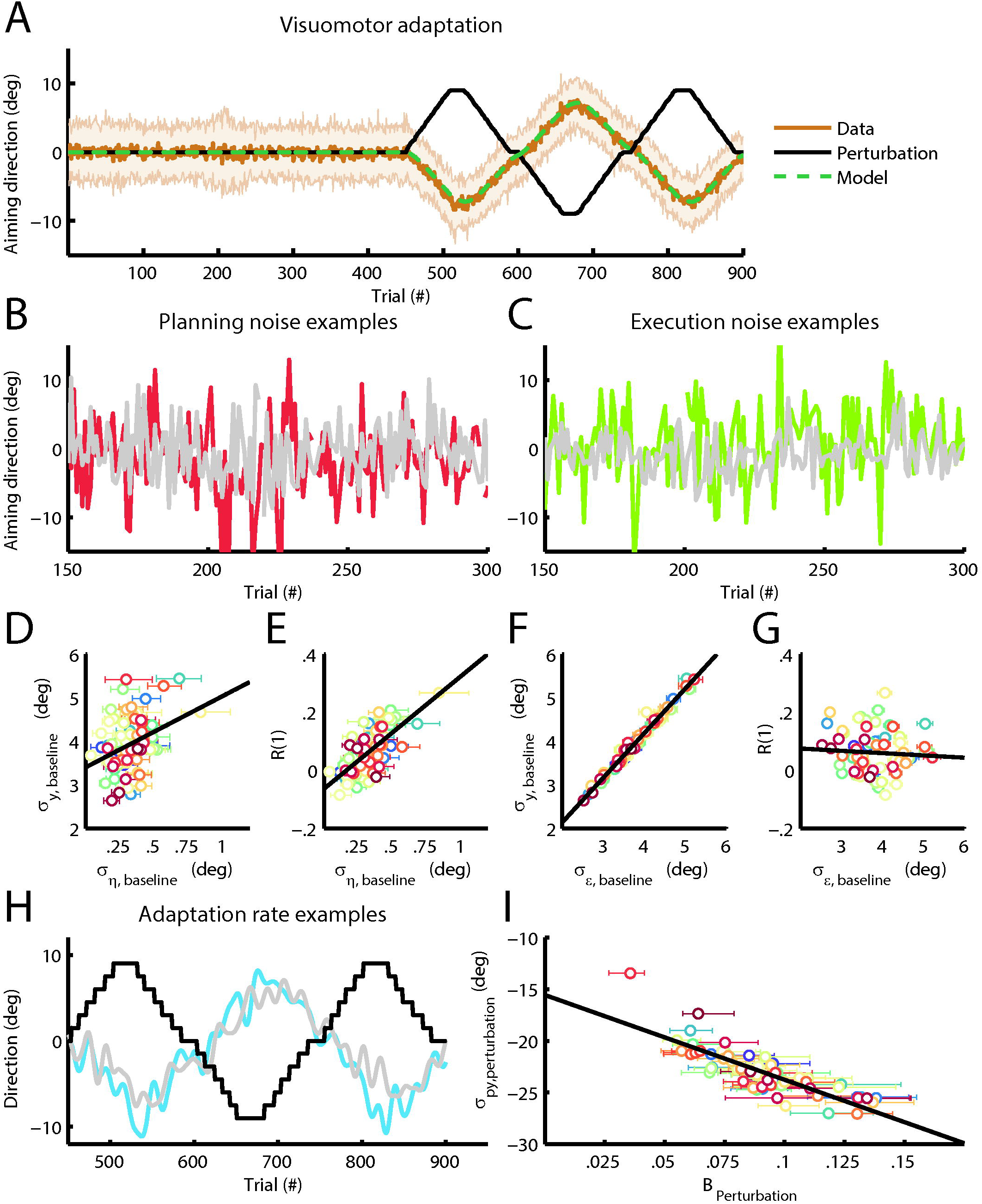
State-space model of visuomotor adaptation. **A.** Visuomotor adaptation. Average aiming traces of the 69 subjects with standard deviations are shown in brown tone colors. The black indicates the average perturbation signal, the green line the model average. **B.** Planning noise examples. The gray line shows a subject with low planning noise (*σ_η,baseline_* = 0.11° *σ_ϵ,baseline_* = 4.0°), the red line a subject with high planning noise (*σ_η,baseline_* = 0.69° *σ_ϵ,baseline_* = 5.0°). **C.** Execution noise examples. The gray line shows a subject with low execution noise (*σ_η,baseline_* = 0.33° *σ_ϵ,baseline_* = 2.7°), the green line a subject with high execution noise (*σ_η,baseline_* = 0.27° *σ_ϵ_* = 5.1°). **D-G** Relation between model estimates and baseline parameters. Models estimates and 68% confidence intervals are shown for every subject as a dot with error bars. The black line is a linear regression between the model estimates and baseline parameters. Panel **D** shows the relation between model estimate *σ_η,baseline_* and baseline parameter *σ_y,baseline_*, panel **E** the relation between model estimate *σ_η,baseline_* and baseline parameter lag-1 autocorrelation *R_Baseline_*(1), panel **F** the relation between model estimate *σ_ϵ,baseline_* and baseline parameter *σ_y,baseline_* and panel **G** the relation between model estimate *σ_ϵ,baseline_* and the baseline parameter *R_Baseline_*(1). **H.** Adaptation rate examples. The thick lines show a slow (gray, *B* = 0.055) and fast subject (blue, *B* = 0.14) smoothened with a 6^th^ order Butterworth filter. The black shows the perturbation signal for the fast subject. **I.** Relation between the model estimate *B_Perturbation_* and perturbation block parameter *σ_py,perturbation_*. Models estimates and 68% confidence intervals are shown for every subject as a dot with error bars. The black line is a linear regression between the model estimates and baseline or perturbation block parameters.

### Parameter validity

Similarly to Cheng and Sabes (2007) we investigated the validity of *B_Perturbation_*[*s*], *σ_η,baseline_*[*s*] and *σ_η,baseline_*[*s*] by correlating the estimates with the variance statistics of the data.

First, standard deviation and lag-1 autocorrelation of the aiming direction are closely linked to planning noise *σ_η_* and execution noise *σ_ϵ_*. Because the baseline set consists of 50% no vision trials, we can neglect the effect of learning term *B*, in which case standard deviation and lag-1 autocorrelation of the aiming direction can be expressed as:

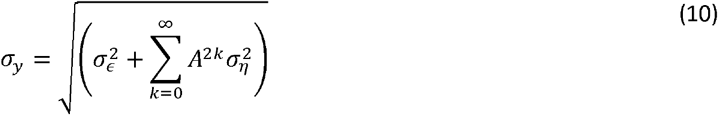

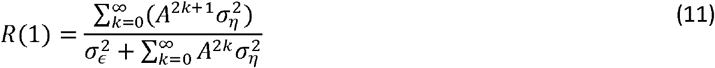

Therefore, standard deviation *σ_y_* increases with both planning noise *σ_η_* (see simulations in Figure 2D) and execution noise *σ_ϵ_* (see simulations in Figure 2F) whereas aiming lag-1 autocorrelation *R*(1) increases with planning noise *σ_η_* (see simulations in Figure 2E) but decreases with execution noise *σ_ϵ_* (see simulations in Figure 2G). Figure 2H shows a simulation of the effect of state and execution noise on aiming direction, and we expect similar relations in the baseline block of our experiment. Figures 3B and 3C show example subjects with low or high baseline planning noise *σ_η,baseline_* (see Figure 3B) and low or high execution noise *σ_ϵ,baseline_* (see Figure 3C). Agreeing with our group level predictions (see Figures 2D-G), we found a positive correlation between planning noise *σ_η,baseline_* and standard deviation *σ_y,baseline_* (*r* = 0.30; 95%HDI = [0.08 0.54]; see Figure 3D), between planning noise *σ_η,baseline_* and aiming lag-1 autocorrelation *R_Baseline_*(1) (*r* = 0.68; 95%HDI = [0.50 0.85]; see Figure 3E) and between *σ_ϵ,baseline_* and standard deviation *σ_y,baseline_* (*r* = 1.00; 95%HDI = [0.96 1.00]; see Figure 3F) and a negligible correlation between *σ_ϵ,baseline_* and aiming lag-1 autocorrelation *R_Baseline_*(1) (*r* = −0.06; 95%HDI = [−0.30 0.17]; see Figure 3G).

Second, the covariance *σ_py_* between the perturbation and aiming direction depends solely on the learning parameters *A* and *B* and is therefore useful to assess the validity of adaptation rate *B* in the perturbation block. The covariance *σ_py_* becomes increasingly negative for higher adaptation rates (see simulations Figure 2I). A simulation of slow and fast learners is given in Figure 2J. We expect a similar relation in the perturbation block of our experiment. Example subjects with a low and high adaptation rate are shown in Figure 3H. Again, according to the model prediction (see Figure 2I), we found a negative correlation between adaptation rate *B_Perturbatlon_* and covariance *σ_py,perturbation_* on a group level (*r* = −0.83; 95%HDI = [−0.97 −0.69]; see Figure 3I).

### Relation between planning noise, execution noise and adaptation rate

We regressed *B_Perturbation_*[*s*] onto *σ_η,baseline_*[*s*] and *σ_ϵ,baseline_*[*s*] and found a positive relation between *σ_η,baseiine_*[*s*] and *B_Perturbation_*[*s*] (*ß* = 0.27 95%HDI=[0.05 0.50]) and a negative relation between *σ_ϵ,baseline_*[*s*] and *B_perturbation_*[*s*] (*ß* = −0.41 95%HDI=[−0.63 −0.16]) with ΔDIC=−9.9 (see Figure 4A-B) in agreement with Kalman filter theory (see Figure 1B-1C).

**Figure 4.**
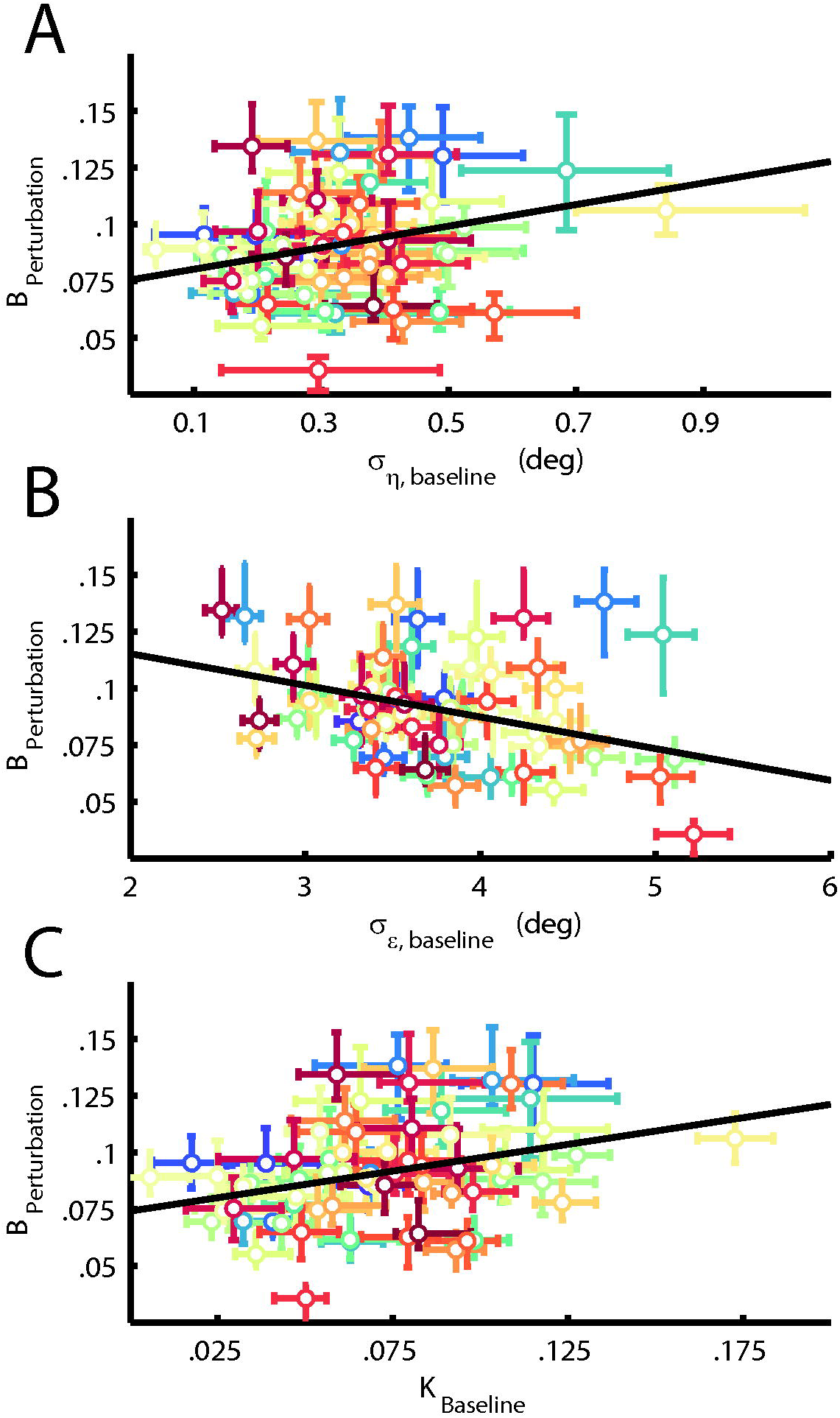
Relation between noise and visuomotor adaptation. **A.** Planning noise and adaptation rate. The black line is a linear regression of *B_Perturbation_* onto *σ_η,baseline_* and *σ_ϵ,baseline_* for average *σ_ϵ,baseline_*. **B.** Execution noise and adaptation rate. The black line is a linear regression of *B_Perturbation_* onto *σ_η,baseline_* and *σ_ϵbaseline_* for average *σ_η,baseline_*. **C.** Kalman gain and adaptation rate. The black line is a linear regression of *B_Perturbation_* onto *K_Baseline_*. Models estimates and 68% confidence intervals are shown for every subject as a dot with error bars.

In addition, we calculated the steady-state Kalman gain for every subject in the baseline blocks from *A*[*s*], *σ_η,baseiine_*[*s*] and *σ_ϵ,baseline_*[*s*] and correlated the steady-state Kalman gain with *B_Perturbation_*. Steady-state Kalman gain was calculated by first solving the Riccati equation for the steady-state covariance *P_∞,baseline_*[*s*]:

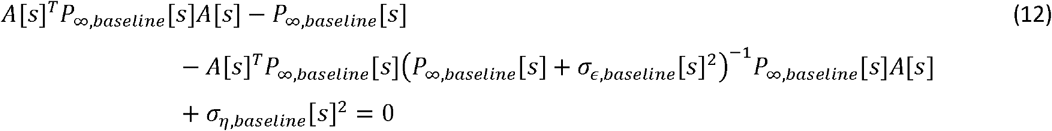

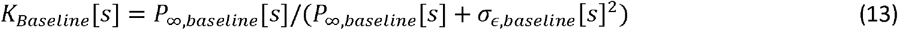

Indeed, we found positive correlations between steady-state Kalman gain *K_Baseline_* and *B_Perturbation_* (*r* = 0.31; 95%HDI = [0.09 0.54]; see Figure 4C).

Finally, we investigated how state and execution noise correlated with baseline aiming peak velocity. Execution noise originates from muscle activity and should increase with vigorous contraction when larger motor units are recruited which fire at a lower frequency and produce more unfused twitches (Harris and Wolpert, 1998; Jones et al., 2002). Indeed, a negligible correlation was found between baseline peak velocity and baseline planning noise *r* = 0.03; 95%HDI=[−0.20 0.25]; whereas a small positive correlation was found between baseline peak velocity and baseline execution noise *r*=0.23; 95%HDI=[0.00 0.47].

#### Control analyses

As control analyses for the fitting procedure, we generated two data sets for our experimental protocol using equations (1)–(4) and the model estimates. In the first dataset, the relation between *B_Block_*[*s*], *σ_η,block_*[*s*] and *σ_ϵ,block_*[*s*] was left unchanged (original dataset), whereas for the second dataset the noise parameters *σ_η,block_*[*s*] and *σ_ϵ,block_*[*s*] were separately permuted in such a way that any regression coefficient between the noise parameters and the adaptation rate would be smaller than 0.05 (permuted dataset). For both datasets, we reestimated the model parameters and expected high test-retest correlations. High test-retest correlations were found for both the ordered (*σ_η,baseline_*= 0.79 [0.64 0.94]; *σ_ϵ,baseline_* = 0.97 [0.93 1.00]; *B_Perturbation_* = 0.89 [0.79 0.99]) and permuted dataset (*σ_η,baseline_*= 0.87 [0.77 0.99]; *σ_ϵ,baseline_* = 0.98 [0.93 1.00]; *B_Perturbation_* = 0.89 [0.78 0.99]). Second, we re-estimated the linear regression of adaptation rate *B_Perturbation_* onto *σ_η,block_* and *σ_η,block_* and expected the correlations to remain for the ordered dataset and disappear for the permuted dataset. Indeed, we found a positive relation between *σ_η,baseline_*[*s*] and *B_Perturbation_*[*s*] (*β* = 0.23 95%HDI=[0.01 0.48]) and a negative relation between *σ_ϵ,baseline_*[*s*] and *B_Perturbation_*[*s*] (*β* = −0.37 95%HDI=[−0.61 −0.15]) with ΔDIC=−8.1 for the original dataset. Furthermore, we found a negligible relation between *σ_η,baseline_*[*s*] and *B_Perturbation_*[*s*] (*β* = −0.05 95%HDI=[−0.31 0.19]) and between *σ_ϵ,baseline_*[*s*] and *B_Perturbation_*[*s*] [*β* = −0.01 95%HDI=[−0.26 0.23]) with ΔDIC=4.0 for the permuted dataset.

## DISCUSSION

We investigated the relation between components of motor noise and visuomotor adaptation rate across individuals. If adaptation approximates optimal learning from movement error, it can be predicted from Kalman filter theory that planning noise correlates positively and execution noise negatively with adaptation rate (Kalman, 1960). To test this hypothesis, we performed a visuomotor adaptation experiment in 69 subjects and extracted planning noise, execution noise and adaptation rate using a state-space model of trial-to-trial behavior. Indeed, we found that adaptation rate in the perturbation block correlates positively with baseline planning noise (r=0.27; 95%HDI=[0.05 0.50]) and negatively with baseline execution noise (r=-0.41; 95%HDI=[−0.63 −0.16]). In addition, the steady-state Kalman gain calculated from baseline state and execution noise correlated positively with adaptation rate in the perturbation block (r = 0.31; 95%HDI = [0.09 0.54]). We discuss implications of our findings for the optimal control model of movement and cerebellar models of adaptation and identify future applications of Bayesian state-space model fitting.

### Optimal control model of movement

The optimal control model of movement has been successful in providing a unified explanation of motor control and motor learning (Todorov and Jordan, 2002). In this framework, the motor system sets a motor goal (possibly in the prefrontal cortex) and judges its value based on expected costs and rewards in the basal ganglia (Shadmehr and Krakauer, 2008). Selected movements are executed in a feedback control loop involving the motor cortex and the muscles which runs on an estimate of the system’s states (Shadmehr and Krakauer, 2008). Both the feedback controller and the state estimator are optimal in a mathematical sense. The feedback controller because it calculates optimal feedback parameters for minimizing motor costs and maximizing performance, given prescribed weighting of these two criteria (Åström and Murray, 2008). The state estimator because it optimally combines sensory predictions from a forward model (cerebellum) with sensory feedback from the periphery (parietal cortex), similar to a Kalman filter (Kalman, 1960; Wolpert et al., 1995). In the optimal control model of movement, motor adaptation is defined as calibrating the forward model, which is optimal in the same sense as the state estimator (Shadmehr et al., 2010).

Wu et al. (2014), is one of the first studies to suggest that there may be a positive relationship between motor noise and motor adaptation. They outlined two apparent challenges of their findings to the optimal control approach: first, they claimed that optimal motor control is inconsistent with a positive relation between motor noise and adaptation rate; second, they claimed that optimal motor control does not account for the possibility that the motor system shapes motor noise to optimize. We take a different view. Because we find that only the planning component correlates positively with adaptation rate, our results are predicted by Kalman filter theory (Kalman, 1960) and consistent with optimal control models of movement (Todorov and Jordan, 2002; Åström and Murray, 2008). However, we do agree that the mathematical structure used to express the optimal control approach does not provide a clear way to discuss shaping noise to optimize adaptation. While this may be a technical difficulty from the point of view of optimal feedback approaches, it is apparent that there is electrophysiological evidence that some animals do shape noise to optimize adaptation. This evidence can be found in monkeys (Mandelblat-Cerf et al., 2009). In addition, studies in Bengalese finches show that a basal ganglia-premotor loop learns a melody from reward (Charlesworth et al., 2012) by injecting noise (Kao et al., 2005) to promote exploration (Tumer and Brainard, 2007) during training (Stepanek and Doupe, 2010) and development (Olveczky et al., 2005). We suggest that a similar mechanism operates in humans during adaptation. This additional tuning mechanism could be an interesting topic of future studies into optimal control models of movement.

### Cerebellar model of motor adaptation

Motor adaptation is the learning process which fine tunes the forward model and is believed to take place in the olivocerebellar system (De Zeeuw et al., 2011). How could this learning process be sensitive to planning noise and execution noise on a neuronal level?

Central to the forward model is the cerebellar Purkinje cell, which responds to selected sensory (Chabrol et al., 2015) and motor (Kelly and Strick, 2003) parallel fiber input with a firing pattern reflecting kinematic properties of upcoming movements (Pasalar et al., 2006; Herzfeld et al., 2015). When Purkinje cell predictions of the upcoming kinematic properties are inaccurate, activity of neurons in the cerebellar nuclei is proportional to the prediction error. This is apparently because inhibitory Purkinje cell input cannot cancel the excitatory input from mossy fibers and the inferior olive (Brooks et al., 2015). The sensory prediction error calculated by the cerebellar nuclei could be used to update either (1) motor commands in a feedback loop with (pre)motor areas (Kelly and Strick, 2003) or (2) state estimates of the limb in the parietal cortex (Grafton et al., 1999; Clower et al., 2001). During adaptation, parallel fiber to Purkinje synapses associated with predictive signals are strengthened and parallel fiber to Purkinje cell synapses associated with non-predictive signals are silenced (Dean et al., 2010). These plasticity mechanisms are affected by climbing fibers originating from the inferior olive, which integrate input from the sensorimotor system and the cerebellar nuclei and act as a teaching signal in the olivocerebellar system (De Zeeuw et al., 1998; Ohmae and Medina, 2015).

No previous experimental or modeling work has considered how planning or execution noise might be conveyed to the cerebellum or how they might influence plasticity. We speculate that planning noise is reflected in synaptic variability of the parallel fiber to Purkinje cell synapse. Electrophysiological studies of CA1 hippocampal neurons have shown that synaptic noise can improve detection of weak signals through stochastic resonance (Stacey and Durand, 2000). Such a mechanism might help form appropriate connections at the parallel fiber to Purkinje cell synapse during adaptation. In addition, theoretical studies on deep learning networks have shown that gradient descent algorithms, which can be likened to error-based learning, benefit from adding noise to the gradient at every training step (Neelakantan et al., 2015). Furthermore, we speculate that execution noise affects adaptation through climbing fiber firing modulation. Execution noise will decrease reliability of sensory prediction errors because (1) the motor plan is not executed faithfully (motor noise) (van Beers et al., 2004) and (2) the sensory feedback is inaccurate (sensory nose) (Osborne et al., 2005). Therefore, when sensory information for a specific movement plan has been unreliable in the past the olivocerebellar system might decrease its response to sensory prediction error, for example by decreasing climbing fiber firing in the inferior olive (De Zeeuw et al., 1998), which would lower the adaptation rate. The existence of such a mechanism has also been suggested by a recent behavioral study which showed a specific decline in adaptation rate for movement perturbations that had been inconsistent in the past (Herzfeld et al., 2014).

### Two-rate models of adaptation

Our results are based on a one-rate learning model of adaptation (Cheng and Sabes, 2006, 2007; van Beers, 2009). However, recent studies have suggested that a two-rate model composed of a slow but retentive and a fast but forgetting learning system provides a better explanation for learning phenomena such as savings and anterograde interference (Smith et al., 2006). Fast learning might take place at the cerebellar cortex and slow learning at other sites such as the cerebellar nuclei or the brain stem, which is supported by electrophysiological studies of vestibulo-ocular reflex adaptation (Blazquez et al., 2004) and eyeblink conditioning (Medina et al., 2001). In addition, an alternative two-rate formulation has been proposed which dissociates an explicit, possibly cortical component which sets the movement goal, and an implicit, possibly subcortical component, which learns from the movement error compared to that movement goal (Mazzoni and Krakauer, 2006; Taylor et al., 2014), although this formulation might overlap with the original two-rate model (McDougle et al., 2015). However, the exact anatomical substrates for these learning mechanisms remain unknown. The experimental design and analysis we used does not allow us to map noise and adaptation rate to learning systems with different rates. Future studies quantifying slow and fast adaptation, or implicit and explicit adaptation could benefit from our Bayesian statistical approach to quantify individual differences in adaptation rate and motor noise.

## Acknowledgements

This work was supported by ZonMw (project # 10-10400-98-008) and Stichting Coolsingel.

